# Genome and epigenome engineering CRISPR toolkit for probing *in vivo cis*-regulatory interactions in the chicken embryo

**DOI:** 10.1101/135525

**Authors:** Ruth M Williams, Upeka Senanayake, Mara Artibani, Gunes Taylor, Daniel Wells, Ahmed Ashour Ahmed, Tatjana Sauka-Spengler

**Affiliations:** University of Oxford, Weatherall Institute of Molecular Medicine, Radcliffe Department of Medicine, Oxford, OX3 9DS, UK; University of Oxford, Ovarian Cancer Cell Laboratory, Weatherall Institute of Molecular Medicine, Oxford, OX3 9DS, UK; Nuffield Department of Obstetrics and Gynaecology, University of Oxford, Women’s Centre, John Radcliffe Hospital, Oxford, OX3 9DU, UK

**Keywords:** CRISPR, Chicken, enhancer, neural crest, dCas9-LSD1, dCas9-KRAB

## Abstract

CRISPR-Cas9 genome engineering has revolutionised all aspects of biological research, with epigenome engineering transforming gene regulation studies. Here, we present a highly efficient toolkit enabling genome and epigenome engineering in the chicken embryo, and demonstrate its utility by probing gene regulatory interactions mediated by neural crest enhancers. First, we optimise efficient guide-RNA expression from novel chick U6-mini-vectors, provide a strategy for rapid somatic gene knockout and establish protocol for evaluation of mutational penetrance by targeted next generation sequencing. We show that CRISPR/Cas9-mediated disruption of transcription factors causes a reduction in their cognate enhancer-driven reporter activity. Next, we assess endogenous enhancer function using both enhancer deletion and nuclease-deficient Cas9 (dCas9) effector fusions to modulate enhancer chromatin landscape, thus providing the first report of epigenome engineering in a developing embryo. Finally, we use the synergistic activation mediator (SAM) system to activate an endogenous target promoter. The novel genome and epigenome engineering toolkit developed here enables manipulation of endogenous gene expression and enhancer activity in chicken embryos, facilitating high-resolution analysis of gene regulatory interactions *in vivo*.

**Summary Statement:** We present an optimised toolkit for efficient genome and epigenome engineering using CRISPR in chicken embryos, with a particular focus on probing gene regulatory interactions during neural crest development.

**List of Abbreviations:** Genome Engineering (GE), Epigenome Engineering (EGE), single guide RNA (sgRNA), Neural Crest (NC), Transcription Factor (TF), Next Generation Sequencing (NGS), somite stage (ss), Hamburger Hamilton (HH).

## Introduction

CRISPR-Cas9 genome editing has been successfully used in a number of model organisms including *Drosophila* (Port et al., 2014), mouse (Wang et al., 2013), *Xenopus* (Guo et al., 2014), zebrafish (Hwang et al., 2013), lamprey (Square et al., 2015) and chick (Veron et al., 2015). CRISPR-Cas9 technology requires two components, a single guide RNA (sgRNA) and a Cas9 endonuclease. The sgRNA includes a user defined target-specific 20bp spacer fused directly to a trans-activating RNA (tracrRNA), which is necessary for efficient Cas9 loading. Cas9 generates a double strand break (DSB), most frequently repaired by the non-homologous end joining (NHEJ) pathway, resulting in disruption of the targeted genomic region by introduction of indels. Only one CRISPR study has been reported in the chicken embryo, whereby the authors employed a tetracycline-inducible Cas9 and Tol2-mediated integration of CRISPR components, to knockout Pax7 at later stages, allowing sufficient time for somatic genome editing (Veron et al., 2015). CRISPR-mediated germ-line editing using cultured primordial germ cells (PGCs) has recently been employed to develop transgenic chicken lines (Dimitrov et al., 2016; Oishi et al., 2016).

Here we have established a highly efficient *in vivo* genome and epigenome engineering toolkit for studying gene regulatory interactions in the early chicken embryo. Our optimised methods not only establish highly penetrant bi-allelic CRISPR-mediated gene knockouts, but also enable use of RNA-guided nuclease-deficient dCas9-effector fusion proteins to directly target and modulate the chromatin landscape and consequently gene expression in a developing embryo. To this end, we have generated and optimised a novel mini-vector system that uses a chick U6 promoter to mediate high, sustained sgRNA expression *in vivo*. We have also created a wild-type Cas9 expression vector with a Citrine reporter, as well as several dCas9-effector constructs that enable epigenome manipulation of endogenous enhancers and promoters.

As a proof of principle, we use this novel toolkit to confirm gene regulatory interactions during early neural crest development. The chicken embryo is an ideal model for probing gene regulatory circuits, as it is amenable to *in vivo* perturbation using highly efficient electroporation methods. Moreover, *ex ovo* bilateral electroporation, where each side of the embryo receives a separate set of plasmids, provides an excellent internal control for each experiment. Our analysis pipeline enabled (i) knockout of upstream transcription factors and assessment of their effects on enhancer activity, (ii) deletion of endogenous enhancers, (iii) epigenetic modulations of endogenous enhancers to assess their role in regulation of the endogenous gene, and (iv) premature activation of endogenous gene loci. The toolkit and optimised protocols developed in this study provide a comprehensive resource to study gene regulatory interactions in the early chicken embryo.

## Results and discussion

### Optimised mini vectors for sgRNA expression

To achieve efficient genome engineering using the CRISPR/Cas9 system, it is essential to maintain high expression of sgRNAs, which are rapidly degraded when not incorporated into Cas9 protein (Hendel et al., 2015). Most current RNA Pol III-dependent systems for expression of sgRNAs in amniotes employ human RNU6-1 promoter, inherited from siRNA expression vectors (Miyagishi and Taira, 2002). However, the optimised *Drosophila* genome engineering toolkit makes use of an alternative Pol III promoter (U6.3), which exhibits much higher activity (Port et al., 2014).

To build an optimal sgRNA expression system in the chicken embryo, we generated four sgRNA mini-vectors, each harbouring a different chick U6 promoter (U6.1, U6.2, U6.3 and U6.4) (Kudo and Sutou, 2005), tracrRNA and BsmBI-flanked cloning cassette (Fig. 1A and Fig. S1A). Using a modified GoldenGate assembly (supp. Protocol 1) we cloned the same spacer targeting the coding region of the FoxD_3_ gene into U6-mini-vectors. We co-electroporated each U6-mini-vector with a plasmid ubiquitously expressing Cas9 (Cas9-2A-Citrine, Fig. S1B) into the entire epiblast of stage HH4 chicken embryos (Hamburger and Hamilton, 1951) and allowed embryos to develop to stage HH10. To measure the efficiency of sgRNA transcription from different U6 promoters, we analysed genome editing events caused by different U6-mini-vector/Cas9 co-electroporations. To this end we dissected cranial dorsal neural tubes of four individual embryos for each U6-mini-vector/Cas9 and Cas9-only controls and assessed the presence of DNA hetero-duplexes within the target region using High Resolution Melt Analysis (HRMA, supp. Protocol 2) (Bassett et al., 2013; Dahlem et al., 2012). U6.1 and U6.3 promoters showed consistent, reproducible evidence of Cas9-mediated mutations, while we observed more variable effects from U6.2 and U6.4 promoters (Fig. 1B). We used the U6.3 promoter-driven sgRNA expression mini-vector (pcU6.3) in all subsequent experiments.

**Figure 1.**
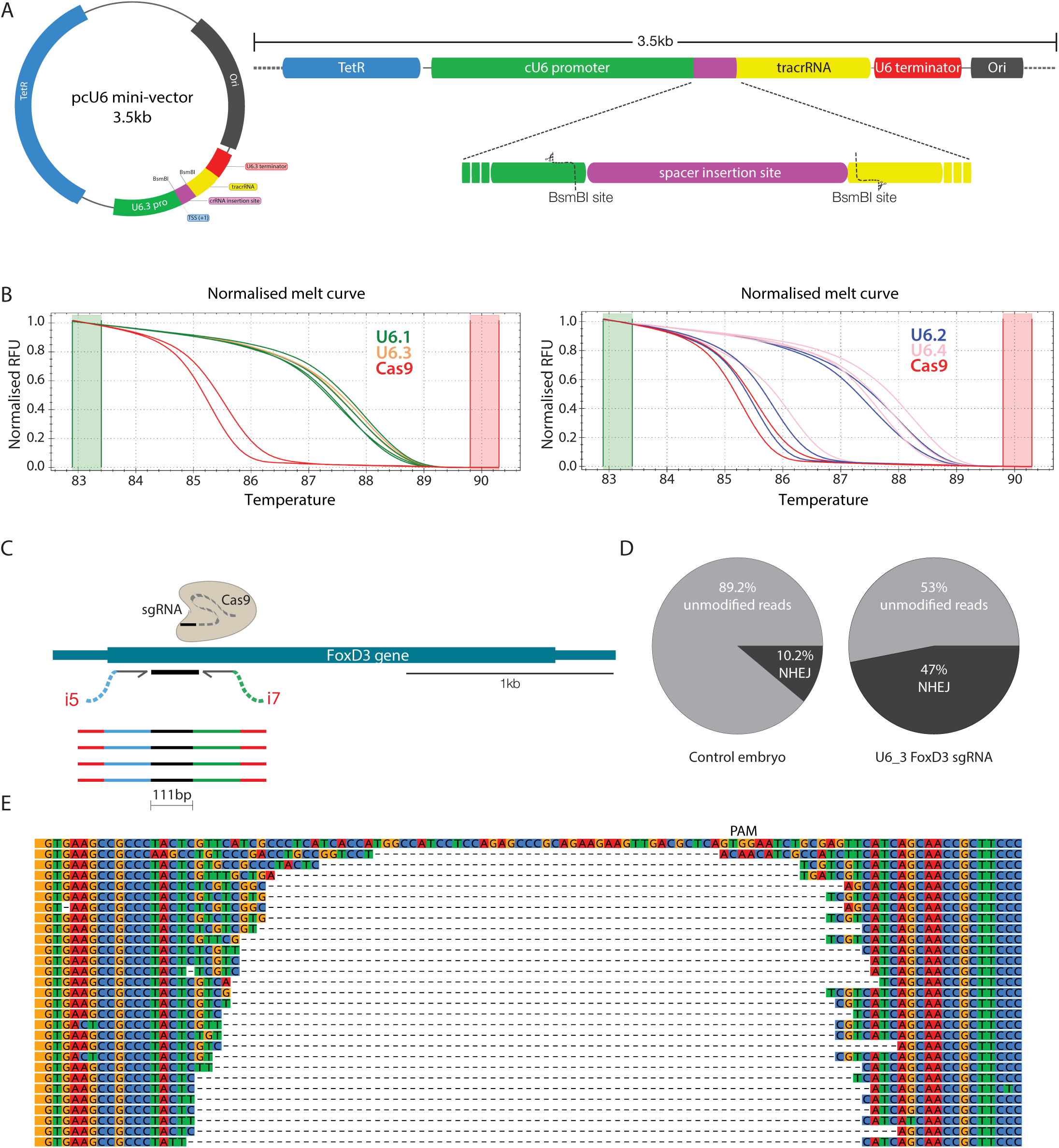
Building sgRNA mini-vectors. (A) 3.5kb sgRNA expression mini-vector (pcU6) contains a chick U6 promoter, BsmBI-flanked sgRNA cassette, tracrRNA scaffold and U6 terminator. (B) Efficacy of four U6 promoters was tested using HRMA. Each HRMA plot shows normalised melt curves from embryos co-electroporated with Cas9 and one of the U6 mini-vectors (pcU6.1, pcU6.2, pcU6.3 and pcU6.4) containing the same sgRNA, and control Cas9-only electroporated embryos. Each HRMA curve is the melting profile of re-annealed amplicons generated from a single embryo, showing temperature shift when compared to Cas9-only controls (red). Relative Fluorescence Units (RFU) for experimental samples were normalised to controls in each case. (C) Schematic of FoxD_3_ genomic structure and targeted NGS approach for testing GE efficiency (D) Pie charts showing the percentage of modified reads for two representative embryos (FoxD_3_ sgRNA-targeted and control Cas9-only). (E) Alignment of a subset of deletion variants identified in the experimental embryos.

To further quantify genome editing efficiency, we profiled CRISPR-mediated somatic indels using a rapid next generation sequencing (NGS) approach. We generated NGS libraries from six individual embryos by amplifying the sgRNA-targeted region with primers that include sequencing adaptors and custom indexes (Fig. 1C; adapted from (Gagnon et al., 2014). Sequencing reads were mapped to the FoxD_3_ amplicon and analysed using the CRISPResso tool (Pinello et al., 2016) to determine the indel frequency. The analysis showed that all experimental embryos had a higher percentage of NHEJ events than the controls (Fig. 1D), as well as multiple deletion variants (Fig. 1E). Controls showed minimal modification as a consequence of five recurrent SNPs.

### Targeting transcription factors controlling enhancers

Having established an efficient sgRNA delivery protocol, we next used CRISPR/Cas9-mediated gene knockout to probe input-enhancer interactions, focusing on targeting transcription factors (TFs) implicated in neural crest development. To this end, we designed sgRNAs targeting Msx1, Pax7, Sox9, c-Myb and Ets1, previously demonstrated to act upstream of the FoxD_3_ enhancer NC1 (Simoes-Costa et al., 2012) or the Sox10 enhancer 10E2 (Betancur et al., 2010) (Fig. 2A). To ensure loss of TF function, sgRNAs were designed to target the essential DNA binding domains encoded within each of these genes (Fig. 2B). sgRNA spacers were predicted manually by scanning regions of interest for proximal PAM sequences, final choices were based on low self-complementarity and unique alignment to the chick genome, cloned into the pcU6.3 mini-vector and tested individually as above (Fig. 2C). In addition to HRMA, we have also adopted an alternative T7 endonuclease assay for candidate sgRNAs validation (Supp. Protocol 3). The selected sgRNAs were co-electroporated with ubiquitous Cas9-2A-Citrine and either FoxD_3_ (NC1) or Sox10 (10E2) enhancer driving mCherry, to assess enhancer reporter activity in the TF knockout condition. We performed bilateral electroporations at HH4, with the left side receiving the target sgRNA+Cas9 and the right side a scrambled sgRNA+Cas9 as an internal control (Fig. 2D). Both sides received the mCherry reporter containing the tested enhancer. Consistent with previous morpholino-based studies (Betancur et al., 2010; Simoes-Costa et al., 2012), we found that knockout of either Msx1, Pax7 or Ets1 strongly reduces mCherry expression mediated by the NC1 enhancer at 5-7 somite stage (ss) in 70%, 100% and 71.4% of cases, respectively (Figs. 2E-G, 2E’-G’; n=10, 7 and 11, respectively). At later stages however, only Pax7 knockouts had a moderate effect (33.3%). Similarly, knockout of Sox9, c-Myb and Ets1 led to a decrease in the activity of Sox10 enhancer, 10E2, with a 100% penetrance for all factors at 5-7ss (Figs. 2H-J, 2H’-J’). However, we observed a decreased effect at later stages (8-10ss; Sox9 14.3% and Ets1 50%), in line with previous findings that Sox10 auto-regulates to maintain 10E2 activity and Sox10 expression (Betancur et al., 2010; Wahlbuhl et al., 2012). These results demonstrate that the CRISPR/Cas9 system is highly efficient in the chicken embryo while re-validating previously shown neural crest-regulatory circuits (Betancur et al., 2010; Simoes-Costa et al., 2012).

**Figure 2.**
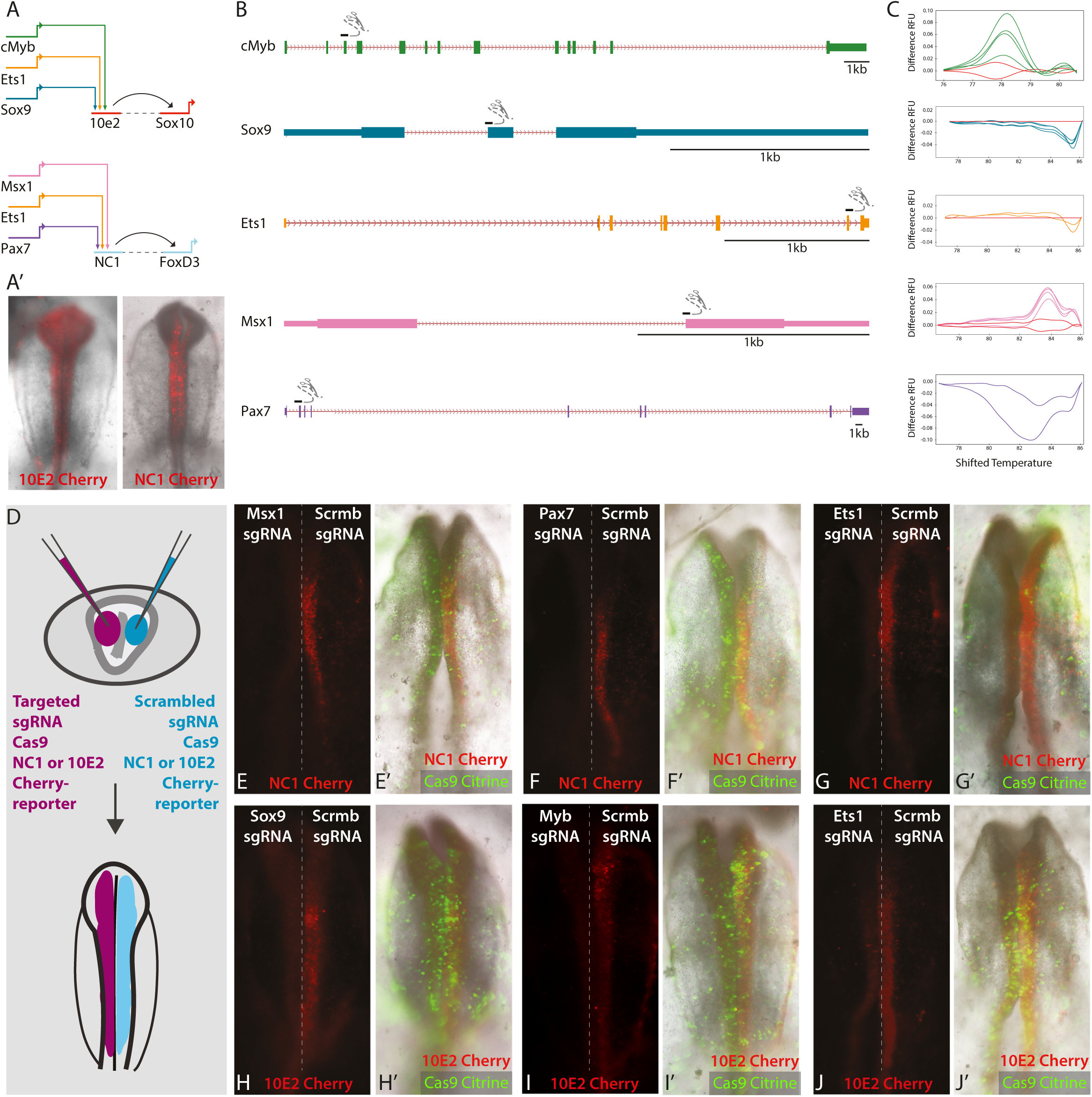
Targeting enhancer inputs reduces enhancer activity. (A) TF inputs into 10E2 and NC1 enhancers. (A’) NC specific mCherry reporter activity controlled by 10E2 and NC1 enhancers. (B) Positions of sgRNAs targeting five TFs. (C) HRMA temperature-shifted difference curves for each sgRNA used. Experimental samples normalised to the controls (Cas9 only, red lines) show significant deviation in each case. (D) Schematic of bilateral electroporation. Target sgRNA, Cas9-2A-Citrine and enhancer-mCherry constructs are co-electroporated on the left (magenta) and scrambled control sgRNA with the same components on the right side of the embryo (blue). (E,F,G) GE-mediated disruption of Msx1, Pax7 or Ets1 results in a decrease in NC1 driven mCherry on the experimental, left side of the embryo (E’,F’,G’) same embryos as in (E, F, G) showing Cas9-2A-Citrine and brightfield view. (H,I,J) GE-mediated disruption of Sox9, cMyb or Ets1 results in a decrease in 10E2-driven mCherry on the experimental side. (H’,I’,J’) same embryos as in (H,I,J) showing Cas9-2A-Citrine and brightfield view. Results confirmed in three independent experiments; representative embryos are shown.

### DSB/NHEJ-mediated removal of targeted enhancers

To fully characterise enhancer function, it is important to study them in their endogenous genomic context. Thus we used our newly developed CRISPR/Cas9 tools to remove the endogenous FoxD_3_-NC1 cranial enhancer (Fig 3A) *in vivo* and assess the consequence of its deletion on the expression of FoxD_3_. We designed and tested sgRNAs flanking the NC1 core region and used these in conjunction with Cas9-2A-Citrine to remove the enhancer. To assay the effect of NC1 knockout, we used bilateral electroporation with target sgRNAs+Cas9 introduced on the left and scrambled control sgRNA+Cas9 on the right side of the same embryo (Fig 3C). Embryos were reared to the desired stages, experimental (left) and control (right) dorsal neural tubes dissected, and the effect on endogenous FoxD_3_ expression was assessed using quantitative RT-PCR (qPCR). We observed a decrease in FoxD_3_ expression on the experimental side in 66.7% of embryos analysed at 5-8ss (n=9) (Fig. 3D). The threshold level of FoxD_3_ expression at its onset (∼0.35 arbitrary units on the control side) was established using the absolute quantification with the same standard curve cDNA across all experiments. This demonstrates that NC1 endogenous enhancer activity is essential for the onset of the FoxD_3_ expression.

**Figure 3.**
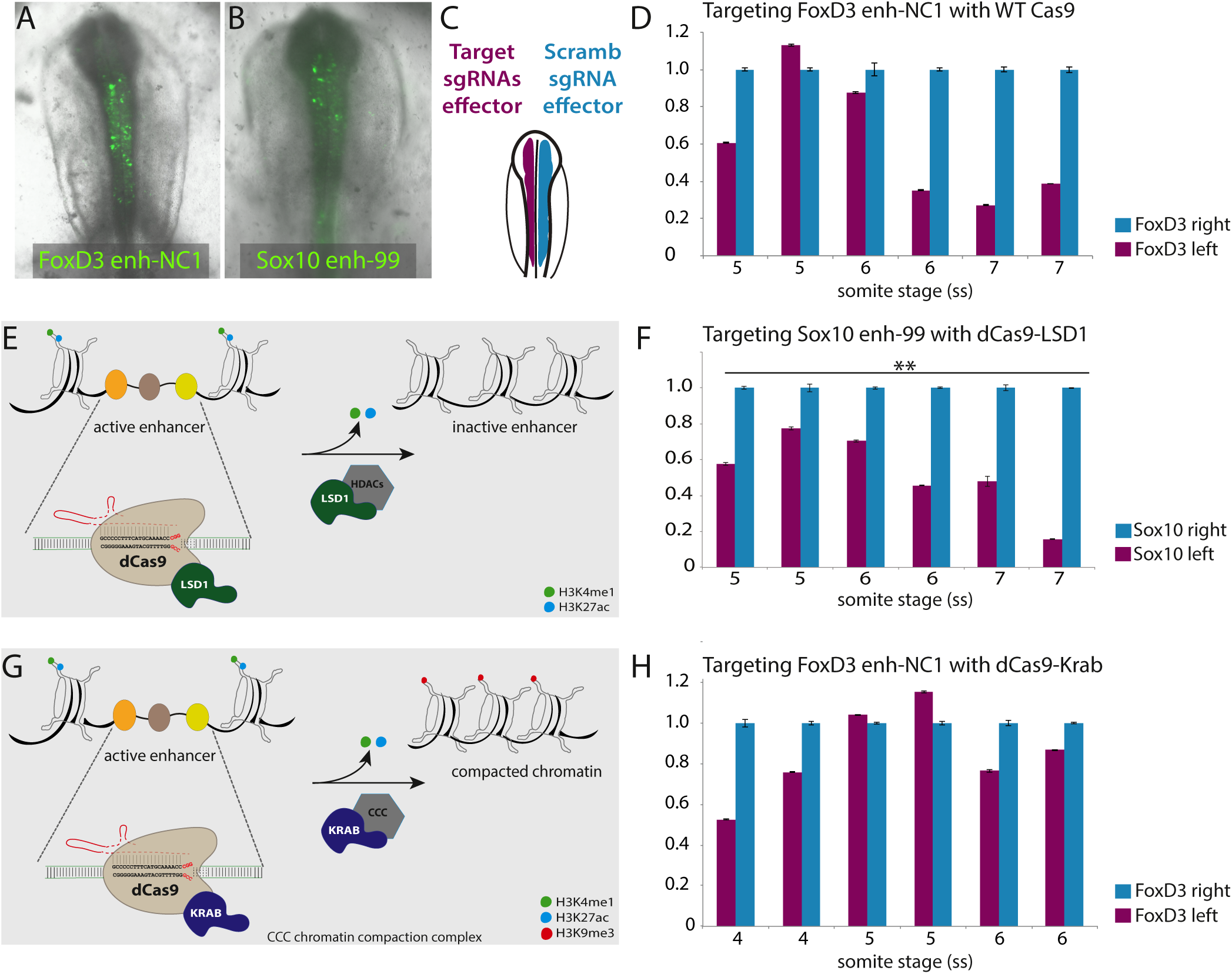
Epigenome engineering modulates endogenous enhancer activity. (A) NC1-driven Citrine expression at 6ss. (B) Novel Sox10 enhancer-99 drives Citrine expression in NC at 7ss. (C) Schematic of bilateral electroporation. (D) Deletion of NC1 using Cas9 and sgRNAs flanking the enhancer core region causes a down-regulation of FoxD_3_ expression on the experimental side (left, magenta) versus control (right, blue) side, (77.7% of embryos, n=16). (E,G) Schematics depict epigenetic enhancer silencing mechanisms by dCas9-LSD1 (E) and dCas9-KRAB (G). (F) 5 guides targeted to the Sox10 enh-99, co-electroporated with dCas9-LSD1 cause a down-regulation of endogenous Sox10 expression at 5-7ss on the experimental compared to the control side (100% of embryos n=9, \*\**p<0.01*). (H) 5 guides targeted to NC1 enhancer, co-electroporated with dCas9-KRAB cause a down-regulation of endogenous FoxD_3_ expression on the experimental versus control side (57% of embryos, n=14). Results confirmed in three independent experiments; representative embryos are shown, error bars represent the standard deviation.

### Epigenomic modification of targeted enhancers

To further refine our analysis of enhancer function *in vivo*, we adopted targeted epigenome engineering (EGE) approaches to alter the chromatin landscape associated with active enhancers. Hitherto, such techniques have only been used *in vitro* (Kearns et al., 2015; Mendenhall et al., 2013; Thakore et al., 2015). To bring EGE methodology into the developing chicken embryo we generated chick expression constructs driving fusion proteins of the catalytically inactive *S. pyogenes* Cas9 (dCas9) with lysine specific demethylase 1 (LSD1) or the Krüppel-associated box (KRAB) domain (Fig. S1C). LSD1 is a lysine-specific demethylase that catalyses the removal of H3K4me1/2 and H3K9me2, which are associated with active and repressive chromatin respectively (Shi et al., 2004) (Fig 3E), whereas the KRAB domain is thought to recruit a chromatin compaction complex thus rendering enhancers inaccessible (Sripathy et al., 2006) (Fig 3G). LSD1 was first used as a TALE-fused effector that demethylates enhancer-associated histone methylation leading to inactivation of targeted enhancers (Mendenhall et al., 2013). More recently, *N. meningitidis* dCas9 fusions with either LSD1 or KRAB were used to inactivate known Oct4 *cis*-regulatory elements in mouse embryonic stem cells (Kearns et al., 2015) and human codon-optimised *S. pyogenes* dCas9-KRAB was used to inactivate HS2 enhancer within the globin locus control region in K562 erythroid leukaemia cells (Thakore et al., 2015).

In order to efficiently repress enhancer activity in chicken embryos, we used five sgRNAs tiled across each targeted enhancer. sgRNA spacers were cloned into the pcU6.3 mini-vector and tested individually, as described above. Pools of five selected sgRNAs per enhancer were co-electroporated bilaterally with the ubiquitous dCas9-effector fusion construct on the left side of the embryo. The right, control side was electroporated with equal molar quantity of scrambled control sgRNA mini-vector combined with the same dCas9-effector (Fig. 3C). Embryos were reared to the desired stages, their cranial regions dissected and the effect of chromatin EGE modification on the targeted enhancer assessed by comparing the levels of endogenous target gene on the experimental and control side using qPCR. The time point for analysis of each tested enhancer was set to approximately three hours before the fluorescent reporter can first be detected, to account for the maturation time of the fluorophore. The dCas9-LSD1 targeted demethylation approach was first applied to repress the activity of the Sox10 enhancer, 10E2. Since no changes were observed in the endogenous level of Sox10 on the experimental versus control side at any of the stages tested (4-8ss) (Fig. S2A), we reasoned that 10E2 element might not be the earliest enhancer controlling Sox10 expression. This assumption is supported by the observation that its fluorescent reporter activity only starts to be detected at ∼8-9 ss. Using epigenomic profiling we have identified a novel Sox10 enhancer (enhancer 99, Fig. 3B), yielding Sox10-like fluorescent reporter activity from 4ss. We hypothesised that this element may be important for the onset of endogenous Sox10 expression. Indeed when dCas9-LSD1 activity was targeted to the enh-99, we achieved a reproducible (100%, n=9, *\*\*p<0.01*) knockdown of endogenous Sox10 expression on the experimental versus control side, in embryos ranging from 5-7ss (Fig. 3F). We next targeted dCas9-KRAB to the FoxD_3_ enhancer NC1, whose fluorescent reporter activity is first detectable at 5ss. When analysed at 4-6ss, we observed a decrease in endogenous FoxD_3_ expression on the experimental side in 57% of embryos (n=14) (Fig. 3H), suggesting that while NC1 is required for proper FoxD_3_ expression, a different, potentially earlier acting enhancer(s) may co-regulate FoxD_3_ onset. Interestingly, targeting dCas9-LSD1 to NC1 enhancer had a weaker effect on FoxD_3_ expression (Fig. S2C), with only 44.4% embryos (n=9) showing FoxD_3_ down-regulation. When we targeted dCas9-KRAB to Sox10 enhancer-99, we observed downregulation of Sox10 in 62.5% of embryos (n=11) (Fig S2D), but only 33.3% of embryos (n=15) showed decrease in Sox10 when 10E2 enhancer was targeted with dCas9-KRAB (Fig S2B). Our results suggest that the two presented EGE approaches have different mechanistic modes of action (Kearns et al., 2015), with one being efficient only at stopping the initiation of the enhancer activity (dCas9-LSD1) while the other (dCas9-KRAB) allowed repression of active elements already engaged in enhancing gene expression. By successfully decommissioning NC specific enhancers *in vivo* we demonstrated their functional importance to the expression of the cognate downstream targets.

### Premature activation of endogenous gene expression *in vivo* using CRISPR-ON

To complement our gene and enhancer loss of function studies, we have adapted the dCas9-VP64-mediated activation of endogenous gene loci *in vivo*. VP64 is a well-established transcriptional activator domain consisting of a tetrameric repeat of the minimal activation domain found in herpes simplex protein VP16 (Seipel et al., 1992). dCas9-VP64 fusion has been successfully used in CRISPR-ON experiments to ectopically activate gene expression (Cheng et al., 2013; Guo et al., 2017). Here, we use a dCas9-VP64 fusion in conjunction with the synergistic activation mediator (SAM) system (Konermann et al., 2015). SAM is a three-component system employing (1) a modified pcU6.3_MS2 vector with a tracrRNA scaffold containing stem loops that associate with bacteriophage MS2 coat protein (MCP), (2) MCP-VP64 and (3) dCas9-VP64 expression construct (Fig. S1A,D). Co-expression of all three components results in saturation of the targeted site with effector molecules, thus enabling a ‘CRISPR-ON’ response using just one sgRNA (Fig. 4A). We tested five sgRNAs to the Sox10 promoter and used the most efficient one to prematurely activate Sox10 expression using the dCas9-VP64/SAM system. Following bilateral electroporation at HH4, we observed an increase in Sox10 expression at 3-4ss on experimental versus control side of the embryo, indicating premature activation of the gene (50% embryos, n=12, Fig. 4B).

**Figure 4.**
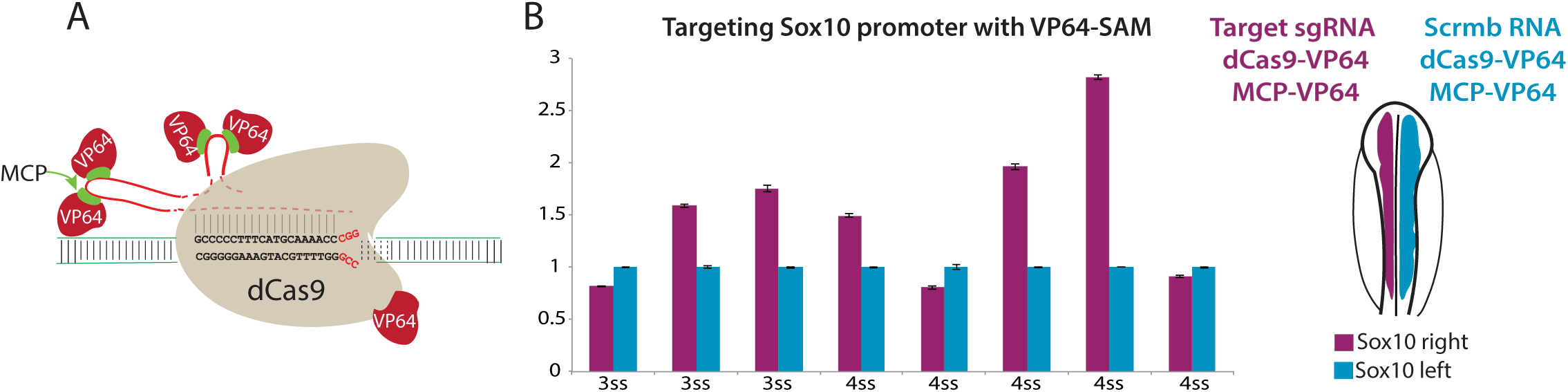
Premature activation of endogenous Sox10 *in vivo* using Synergistic Activation Mediator (SAM). (A) Schematic of dCas9-VP64-MCP SAM complex. (B) One sgRNA targeting the Sox10 promoter co-electroporated with dCas9-VP64 and MCP-VP64 causes a premature activation of Sox10 transcription on the experimental (left, magenta) as compared to the control side (right, blue), 50% embryos, n=12, *\*p<0.05*). Results confirmed in three independent experiments; representative embryos are shown, error bars represent the standard deviation.

Here, we have optimised CRISPR/Cas9 approaches to perform genome and epigenome engineering in the early chicken embryo, with a particular focus on probing gene regulatory interactions. The presented toolkit enables deletion, repression and activation of endogenous enhancers and promoters, thus extending the epigenome engineering approaches, previously only attempted *in vitro* (Kearns et al., 2015; Konermann et al., 2015; Mendenhall et al., 2013; Thakore et al., 2015) to an in vivo context of the developing embryo. Although we focus on the early chicken embryo due to its accessibility for experimental manipulation via electroporation, we propose that such approaches could be easily adapted to other amniotes.

## Materials and Methods

### Construct cloning

Chick U6 sgRNA expression mini-vectors were cloned by replacing the BsaI-flanked cassette from pNG1-pNG4 vector backbones (Cermak et al., 2011) (Addgene #30985-8) with custom-synthesised gBlocks (IDT) containing chick cU6_1-4 promoters (Kudo and Sutou, 2005).

pCAG_Cas9-2A-Citrine construct was generated by removing IRES-H2B-RFP cassette from pCI_H2B-RFP vector (Betancur et al., 2010) and inserting Cas9-2A-Citrine fragment by In-fusion HD (Clontech, cat #638910) cloning.

To construct the pX330 dCas9-LSD1 vector, Cas9m4-VP64 was amplified from (Addgene #47319) and cloned into pX330 (Addgene #42230). VP64 was then removed by EcoRI digest and LSD1 was inserted from EMM67 (Addgene #49043) (Mendenhall et al., 2013). pX330 dCas9-KRAB was generated by In-fusion of a synthetic gBlock containing the KRAB sequence into the EcoRI-linearised pX330-dCas9-LSD vector.

pCAG-dCas9-KRAB-2A-GFP was generated by amplifying dCas9-KRAB-2A-GFP sequence from pLV hUbC dCas9 KRAB T2A GFP (Addgene #71237) and cloning into the pCI-H2B-RFP vector linearised with NotI and XhoI.

The MCP-VP64 construct was generated by removing the IRES-H2B-RFP cassette from pCI_H2B-RFP vector (Betancur et al., 2010) and inserting the MCP and VP64 fragments (amplified from Addgene #61423 and #47319, respectively) using 2-fragment Infusion cloning.

### Embryo culture and electroporations

Fertilised wild type chicken eggs were obtained from Henry Stewart & Co (Norfolk). Staged according to (Hamburger and Hamilton, 1951) and electroporated as previously described (Sauka-Spengler and Barembaum, 2008; Simoes-Costa et al., 2012). Electroporation conditions are detailed in a separate protocol (Supplemental Info).

### GE reagent cloning and validation

Detailed standardised protocols describing guide RNA selection, GoldenGate cloning, bilateral electroporation and validation by HRMA, T7 assays and NGS are available as Supplemental Info and at our resource page (http://www.tsslab.co.uk/resources). All plasmids are available from Addgene (https://www.addgene.org/Tatjana_Sauka-Spengler/).

### qPCR analysis of endogenous levels

RNA extractions of dissected embryonic tissue were carried out using the RNAqueous^®^-Micro Kit (Life Technologies, cat # AM1931). Oligo-dT-primed cDNA was synthesised using Superscript III reverse transcriptase (Invitrogen) and qPCRs performed using Fast SYBR Green reagent (ThermoFisher, cat #4385612) on the Applied Biosystems 7500 Fast Real-Time PCR System. Standard curve method was used to quantify the gene expression. In all embryos, contralateral side was used as an internal control. Statistical significance of bilateral electroporations was determined by generating a control embryo group that received scrambled sgRNAs on both sides and calculating p-values using chi-squared statistical test, based on contingency tables comparing control and experimental embryo groups. *p<0.05, **p<0.01.

### Imaging analysis

Embryos were imaged on an Olympus MVX10 stereomicroscope with 2-2.5x objective using Axio Vision 4.8 software.

## Acknowledgements

We thank Prof Tudor Fulga for helpful advice on the project and the manuscript and Francesco Camera for technical assistance.

### Competing interests

The authors declare no competing or financial interests.

### Author contributions

US, RW and TSS conceived the study. US generated the constructs and RW performed majority of the GE and EGE experiments. MA generated dCas9-VP64/SAM constructs and established protocols for targeted NGS validation of GE events. GT optimised sgRNA GoldenGate cloning and built dCas9-KRAB constructs. DW helped build and test U6 mini-vectors and developed methods for multiple sgRNA expression; RW, MA and TSS analysed the data; RW and TSS wrote the manuscript; RW, US, MA, GT, AA, TSS edited the manuscript; TSS supervised the study.

### Funding

This study was funded by MRC (G0902418), OCA (MA), Lister Institute summer studentship to DW.

